# Multiplex PCR Assay for Clade-typing *Salmonella* Enteritidis

**DOI:** 10.1101/2022.08.12.503823

**Authors:** Sarah Gallichan, Blanca M. Perez-Sepulveda, Nicholas A. Feasey, Jay C. D. Hinton, Juno Thomas, Anthony Marius Smith

**Affiliations:** Centre for Enteric Diseases at the National Institute for Communicable Diseases (NICD), Johannesburg, South Africa; Department of Clinical Microbiology and Infectious Diseases, School of Pathology, Faculty of Health Sciences, University of the Witwatersrand, Johannesburg, South Africa; Department of Clinical Sciences, Liverpool School of Tropical Medicine, Pembroke Place, Liverpool, UK; Malawi Liverpool Wellcome Research Programme, Kamuzu University of Health Sciences, Blantyre, Malawi; Clinical Infection, Microbiology & Immunology, Institute of Infection, Veterinary & Ecological Sciences (IVES), University of Liverpool, Liverpool, United Kingdom

**Keywords:** Non-typhoidal *Salmonella*, real-time PCR, phylogeny, molecular surveillance

## Abstract

2.

*Salmonella* Enteritidis is one of the most commonly reported serovars of non-typhoidal *Salmonella* causing human disease and is responsible for both gastroenteritis and invasive non-typhoidal *Salmonella* (iNTS) disease worldwide. Whole-genome sequence (WGS) comparison of *Salmonella* Enteritidis isolates from across the world have identified three distinct clades, named Global Epidemic, Central/East African and West African, all of which have been implicated in epidemics: the Global Epidemic clade was linked to poultry-associated gastroenteritis, while the two African clades were related to iNTS disease. However, the distribution and epidemiology of these clades across Africa is poorly understood because identification of these clades currently requires whole genome sequencing capacity. Here, we report a sensitive, time- and cost-effective real-time PCR assay capable of differentiating between the *Salmonella* Enteritidis clades to facilitate surveillance and to inform public health responses.

**Impact statement:** Challenges in the diagnosis and treatment of invasive *Salmonella* Enteritidis (*S*. Enteritidis) bloodstream infections in sub–Saharan Africa are responsible for a case fatality rate of approximately 15% (12). It is important to identify distinct clades of *S*. Enteritidis in diagnostic laboratories in the African setting to determine whether particular outbreaks are associated with different health outcomes. Here, we have described the development of a high-quality molecular classification assay for the clade-typing of *S*. Enteritidis that is ideal for use in public health laboratories in resource-limited settings.

## 4. Introduction

The key human pathogen *Salmonella enterica* has over 2,500 serovariants, determined by O and K surface antigens, and within individual serovars, there can be distinct pathotypes that currently require whole genome sequencing (WGS) to identify. This diversity makes it challenging for surveillance systems to identify lineages of concern and thus the importance of specific variants is not communicated to public health authorities and policy makers. Sub-Saharan African (sSA) countries bear the greatest global burden of foodborne disease, are under pressure to increase production of protein-rich foods, often in the form of meat, but often have limited food, water and environmental surveillance capacity (1).

The best available evidence suggests that animal-source foods are the primary origin of foodborne pathogens in sSA (2). With global poultry production surpassing pork production in 2018, it is understandable that the poultry-associated non-typhoidal *Salmonella* (NTS) serovar, *Salmonella* Enteritidis, (*S*. Enteritidis), is the most reported foodborne pathogen in sSA (3). Generally, *S*. Enteritidis infections are associated with outbreaks of gastroenteritis in Europe and the United States (4,5). However, *S*. Enteritidis infections in sSA regions are commonly associated with severe, invasive bloodstream infections, known as invasive non-typhoidal Salmonella disease (hereafter named iNTS) (6,7).

The disproportionately high number of iNTS infections in sSA - approximately 79% of the global burden of iNTS (a 2017 estimate) – is closely associated with the high-risk populations in sSA (high numbers of advanced HIV infections, malaria and young children with immature immune systems) (7). The high prevalence of immunosuppressed individuals in sSA has facilitated the emergence of iNTS as a major public health problem, with the two key serovars being *S*. Enteritidis and *S*. Typhimurium (8,9) A 2016 study investigated the diversity of *S*. Enteritidis in sSA and, in addition to a globally prevalent poultry associated lineage, identified two geographically distinct groups of *S*. Enteritidis strains circulating in sSA, namely the West African and the Central/ Eastern African clade (hereafter the “East African clade”) (8). The West and East African clades were quite distinct from *S*. Enteritidis strains commonly associated with global gastroenteritis outbreaks, the Global Epidemic clade, raising the possibility of different ecological niche adaptation (8).

Despite the recognition of distinct *S*. Enteritidis clades and the severity of iNTS disease, the distribution and epidemiology of these clades across sSA remains poorly understood (10,11). The lack of data pertaining to *S*. Enteritidis clades in sSA is, in part, due to the lack of a distinct molecular typing system for *S*. Enteritidis (12–14). The closely related S. Typhimurium has similarly unique clinical and epidemiological characteristics between S. Typhimurium subtypes that can be clustered using Multi-Locus Sequence Typing (MLST). Indeed, sequence type (ST) 313 has been associated with epidemics of bloodstream infection, in contrast with the globally distributed ST19 that is mostly associated with gastroenteritis (15,16). However, MLST fails to distinguish between S. Enteritidis variants, with the majority of the isolates being assigned to ST11 (17). This becomes epidemiologically problematic when outbreaks of pathologically distinct S. Enteritidis clades are treated as a singular sequence type.

For public health officials and policy makers to both be aware of iNTS as a cause of severe febrile illness and institute policy to interrupt transmission and prevent iNTS, there needs to be capacity to make the distinction between the gastroenteritis-associated Global clade and the multidrug-resistant, invasive infection-associated East and West African clades (12). Currently, the best way to distinguish between S. Enteritidis clades is through whole-genome sequencing, which is not widely available in sSA (19). Ideally, regional public health laboratories need access to robust, accurate and cost-effective tests with a rapid turnaround time capable of differentiating between genetically similar isolates in order to facilitate appropriate epidemiological investigation of distinct pathovariants.

Real-time PCR assays are a commonly used method for the highly specific and sensitive classification of foodborne diseases, thus are widely available (20). When the real-time PCR assay is multiplexed, it has the advantage of being able to identify multiple pathogens with a single assay (20). The scalability and rapid turn-around-time of real-time PCR assays is also beneficial for use in diagnostic settings (20). The aim of the real-time PCR assay developed in this study is to classify *S*. Enteritidis isolates into clades in order to assist laboratories in typing *S*. Enteritidis strains to aid in surveillance of variants with an identical antigenic formula, but which require different public health responses.

## 5. Methods

In respect of the phyletic structure of *S*. Enteritidis, we have designed primers to distinguish three clades and an outlier cluster in a single reaction. These are henceforth denoted “Regional” and “Clade”. The purpose of the Regional (African or Global classification) and Clade (Global Epidemic, Global Outlier, East or West African classification) assays is to further classify *S*. Enteritidis isolates to better understand the transmission and epidemiology of each *S*. Enteritidis clade. The Regional and Clade assays described here are limited to previously confirmed *S*. Enteritidis isolates.

### Control panel isolates

The control panel consisted of 12 *S*. Enteritidis strains that were used as positive controls in the development of the multiplexed real-time PCR assays. The 12 *S*. Enteritidis isolates were obtained as part of the 10,000 *Salmonella* Genomes project (21) and were selected based on the previously published *S*. Enteritidis global population structure predicted by the hierBAPS (hierarchical Bayesian Analysis of Population Structure) algorithm (10). The control panel was assembled to represent the East African (n=3), West African (n=3), Global Epidemic (n=3) and Global Outlier (n=3) clades (Table 1). The clades were grouped according to region, the Global (Global Epidemic and Global Outlier) and African (East African and West African) regions (Table 1). All *S*. Enteritidis samples were stored at −70°C in 500 μL Tryptic Soy broth media (1 L distilled water, 17 g casein, 5 g NaCl, 3 g soytone, 2.5 g dextrose, 2.5 g dipotassium phosphate, adjusted to pH 7.3).

**Table 1.**
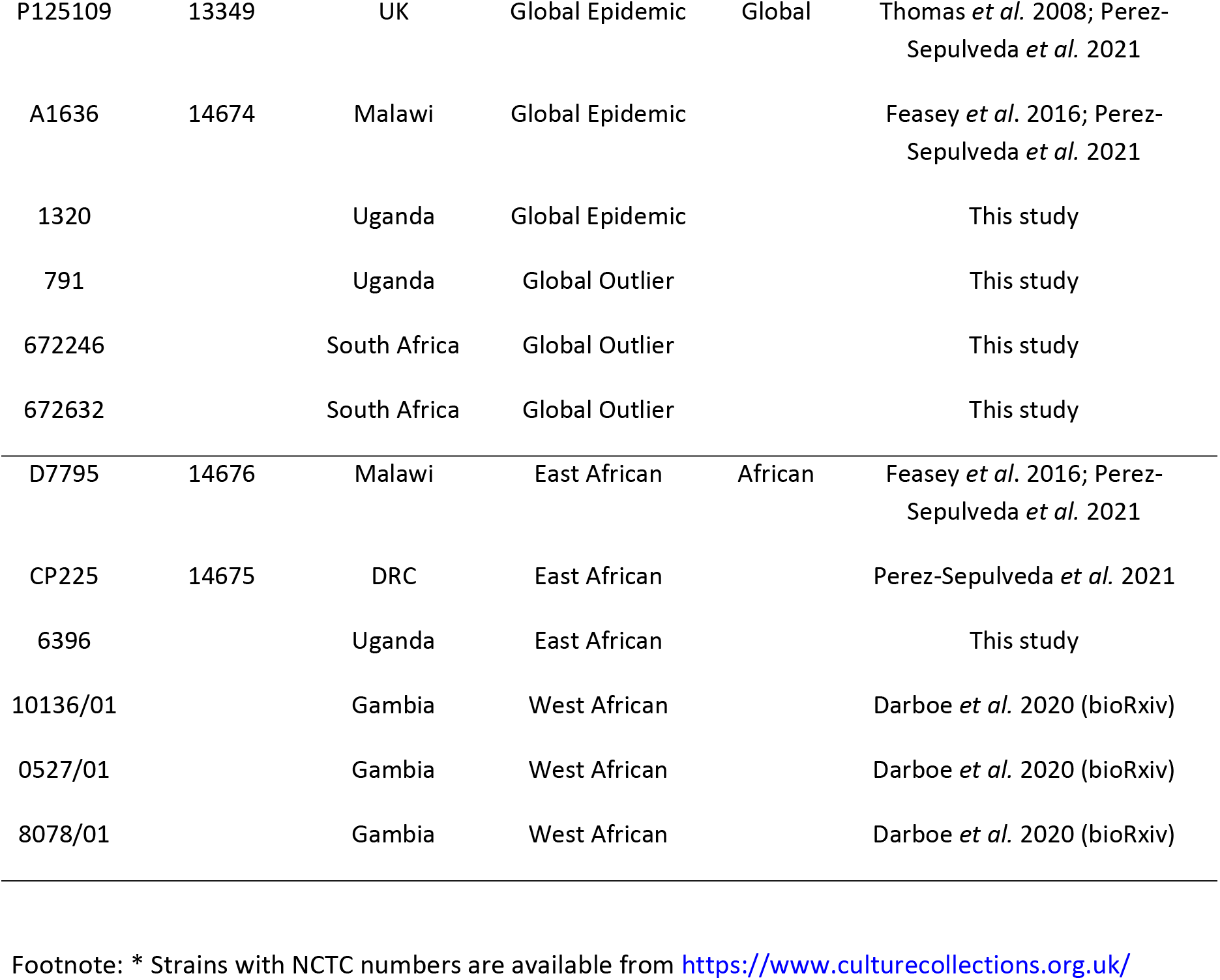
Salmonella Enteritidis strains used as a control panel in this study

| Isolate name | NCTC number* | Origin | Clade | Cluster | References |
| --- | --- | --- | --- | --- | --- |
| P125109 | 13349 | UK | Global Epidemic | Global | Thomas *et al.* 2008; Perez-Sepulveda *et al.* 2021 |
| A1636 | 14674 | Malawi | Global Epidemic |  | Feasey *et al.* 2016; Perez-Sepulveda *et al.* 2021 |
| 1320 |  | Uganda | Global Epidemic |  | This study |
| 791 |  | Uganda | Global Outlier |  | This study |
| 672246 |  | South Africa | Global Outlier |  | This study |
| 672632 |  | South Africa | Global Outlier |  | This study |
| D7795 | 14676 | Malawi | East African | African | Feasey *et al.* 2016; Perez-Sepulveda *et al.* 2021 |
| CP225 | 14675 | DRC | East African |  | Perez-Sepulveda *et al.* 2021 |
| 6396 |  | Uganda | East African |  | This study |
| 10136/01 |  | Gambia | West African |  | Darboe *et al.* 2020 (bioRxiv) |
| 0527/01 |  | Gambia | West African |  | Darboe *et al.* 2020 (bioRxiv) |
| 8078/01 |  | Gambia | West African |  | Darboe *et al.* 2020 (bioRxiv) |

Footnote: \* Strains with NCTC numbers are available from https://www.culturecollections.org.uk/

### Genomic DNA extraction

The control panel isolates were streaked on 5% blood agar (Diagnostic Media Products, Johannesburg, South Africa) plates and incubated overnight in an IN 750 incubator (Memmert, Schwabach, Germany) at 37°C. Single colonies were resuspended in 400 μL of 10X TE buffer (800 mL distilled water, 2.92 g Tris, 15.76 g EDTA (pH 8)) in 2 mL Safe-Lock tubes (Eppendorf, Hamburg, Germany). The QIAamp DNA Mini Kit (QIAGEN, Hilden, Germany) was used to extract genomic DNA according to the instructions provided by the manufacturer. Final DNA concentrations were quantified fluorometrically using the Qubit 2.0 Fluorometer (ThermoFisher Scientific, California, USA).

### Whole-genome sequencing

The control panel isolates were sequenced and assembled as part of the 10,000 *Salmonella* Genomes project using the LITE pipeline for library construction, and Illumina HiSeq™ 4000 system (Illumina, California, USA) (21). Whole-genome sequences of all 12 *S*. Enteritidis isolates were annotated using Prokka v.1.14.5 (22). The resulting annotated genomes were analysed using ROARY v.3.11.2 (23), producing a gene presence/ absence matrix that compared the gene differences across the whole genome of each of the control panel isolates.

### Development of the multiplexed real-time PCR assays

Target genes for the clade-typing real-time PCR assay were selected based on the presence/ absence matrix (Figure 1). To confirm the specificity of the selected genes, a clade typing of 500 of the *S*. Enteritidis genomes available on EnteroBase v1.1.3 was performed. A “Workspace” with the 500 of the *S*. Enteritidis genomes used in the published *S*. Enteritidis global population analysis (10) from whole genome sequences obtained as part of the 10,000 *Salmonella* Genomes project (21) was created. A custom multi-locus sequence typing analysis scheme using the target genes from the clade-typing real-time PCR assay was then used to type the *S*. Enteritidis genomes into clades. The clade results from this EnteroBase query were then compared with the *S*. Enteritidis global population structure predicted by the hierBAPS algorithm (8).

**Figure 1.**
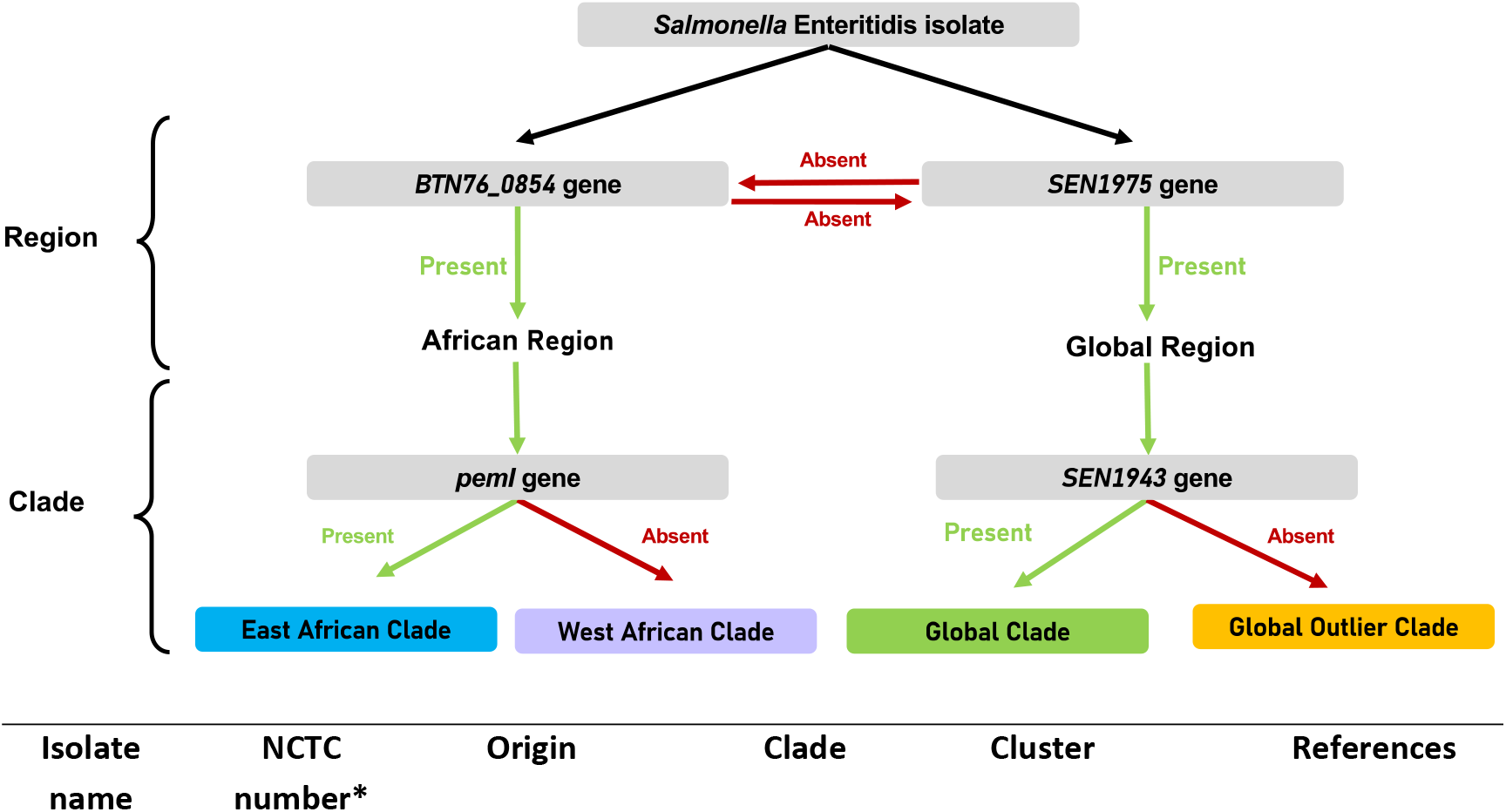
Workflow depicting the clade-typing of a *Salmonella* Enteritidis isolate based on the presence (green) or absence (red) of genes targeted by the clade-typing real-time PCR assay

### Real-time PCR assay conditions

All primers and probes were diluted to a concentration of 20 μM using nuclease-free water (Ambion, ThermoFisher Scientific, California, USA). Four master mixes for the two multiplex real-time PCR assays (Regional assay and Clade assay) were prepared as summarized in Table 2. A real-time PCR assay was set up using 25 μL of TaqMan Gene Expression Master Mix (ThermoFisher Scientific, California, USA), 17.8 μL of nuclease-free water (Ambion, ThermoFisher Scientific, California, USA), 3 μL of the relevant Master Mix (Table 2) Master Mix One for the Regional assay and Master Mix Two for the Clade assay) and 1.2 μL of DNA template to each well of the MicroAmp^®^ Optical 96-well reaction plate (Applied Biosystems, ThermoFisher Scientific, California, USA). In each run, a negative control (1.2 μL of nuclease-free water instead of DNA template) was added to the last well of the MicroAmp® Optical 96-well reaction plate. The wells were then sealed with a MicroAmp® Optical Adhesive Film (Applied Biosystems, Life Technologies™, California, USA) and centrifuged at 15 000 RPM for 1 minute using an AllegraTM X-22R Centriuge (Beckman Coulter™, California, USA) to ensure all reagents were concentrated at the bottom of the wells. The plate was then loaded into a 7500 Real Time PCR System (Applied Biosystems, Life Technologies™, California, USA) and set up with the 7500 Real Time PCR System software version 2.0 (Applied Biosystems, Life Technologies™, California, USA). The reactions underwent PCR amplification as follows: 50°C for 2 min, followed by 95°C for 10 min and 40 cycles of 95°C for 15 s, 60°C for 30 s and 72°C for 30 s.

**Table 2.** Constituents of the two Master Mixes used in the Regional Master Mix and Clade Master Mix real-time PCR

|  | Regional Master Mix |  | Clade Master Mix |  |
| --- | --- | --- | --- | --- |
|  | African | Global | Global Epidemic clade | East African clade |
| Forward primer | African-F | Global-F | Epidemic-F | East-F |
| Reverse primer | African-R | Global-R | Epidemic-R | East-R |
| Probe | African-FAM | Global-CY5 | Epidemic-CY2 | East-FAM |

### Multiplex RT-PCR assay performance

To determine the efficiency of the multiplex real-time PCR assay, 10-fold serial dilutions of genomic DNA extracted from two control isolates (D7795 and A1636) were prepared. The DNA concentration of each dilution was quantified spectroscopically using a NanoDrop 1000 Spectrophotometer (ThermoFisher Scientific, California, USA). A real-time PCR assay was then set up as described above using Master Mix One and Two for the Regional assay and Three and Four for the Clade assay. The DNA concentration yielding the highest Ct value below 30 cycles was determined to be the limit of detection for that primer and probe set, in three technical replicates. The linear range (R^2^) was calculated for the Ct values of the triplicate assays for each primer and probe set using the CORREL function in Microsoft Excel 2010. The slopes of calibration curves were used to calculate the amplification efficiency (PCR efficiency = 10^-1/slope^ – 1) (24).

## 6. Results and discussion

### Oligonucleotide design

A gene presence/absence matrix produced from the pan genome analysis of the 12 control panel isolates was used to identify unique gene target sequences that distinguished clades associated with different geographical regions. These included the *BTN76_08545* gene (family protein accession number WP_023229131.1) for the African region and the *SEN1975* gene for the Global region (family protein accession number WP_001075993.1). Individual clades were recognised using the *SEN1943* gene (family protein accession number WP_058658682.1) for the Global Epidemic clade and *pemI* (protein accession number WP_096198836.1) gene for the East African clade. To determine the sensitivity of the selected genes, a multi-locus query based on the presence/absence of genes *BTN76_08545, SEN1975, SEN1943* and *pemI* in the whole-genome sequences from 500 *S*. Enteritidis isolates was performed using EnteroBase v 1.1.3. When compared with the clade outcome predicted by the hierBAPS algorithm on the 500 *S*. Enteritidis whole-genome sequences (10), the multi-locus classification was 90% effective in predicting the clade of the *S*. Enteritidis isolate (Supplementary Table 1).

The four selected genes were then used to design primers and probes using the online PrimerQuest tool (Integrated DNA Technology; accessible online: https://eu.idtdna.com/pages/tools/primerquest) (Table 3). The specificity of the designed primers and probes was tested on the whole-genome sequences of the 12 control panel isolates using the *in silico* PCR tool in CLC Genomics Workbench version 11.0.1 (QIAGEN, Hilden, Germany). The African cluster primer set amplified an 82 bp fragment of the *BTN76_08545* gene for all six African isolates tested (isolates 10136/01, 0527/01, 8078/01, D7795, CP255, and 6396). The Global region primer set amplified a 126 bp fragment of the *SEN1975* gene for all six Global region isolate sequences (isolates P125109, A1636, 1320, 791, 672246, and 672632). The East African clade primer set amplified a 101 bp fragment of the *pemI* gene from the East African clade isolate sequences (isolates D7795, CP255, and 6396). The Global Epidemic clade primer set amplified an 85 bp fragment of the *SEN1943* gene from the Global Epidemic clade isolate sequences (isolates P125109, A1636, and 1320) (Supplementary Table 2).

**Table 3.**
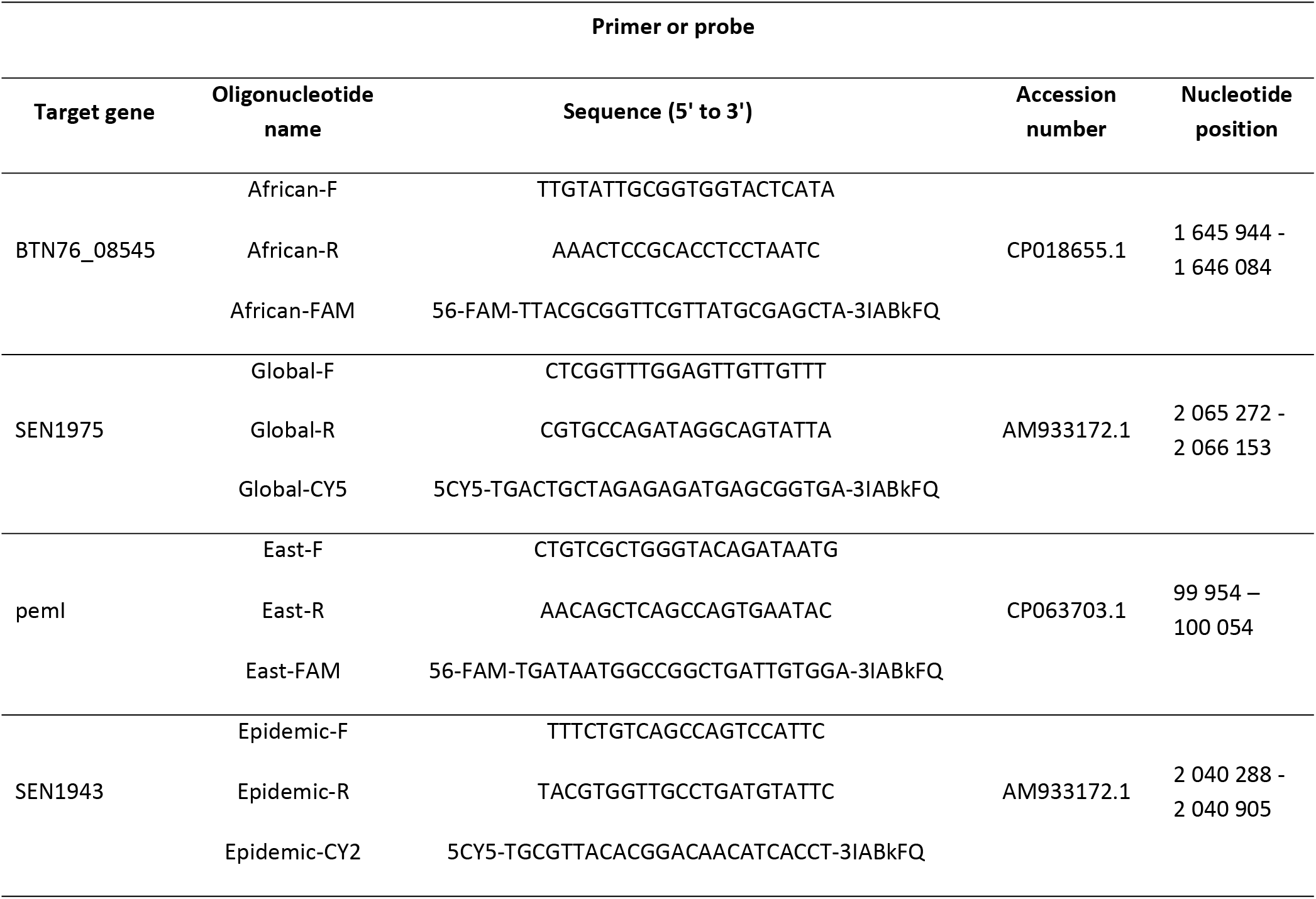
Primer and probe sequences for the development of the *Salmonella* Enteritidis clade-typing real-time PCR assay

### Validation of the real-time PCR assays

The clade-typing real-time PCR assay strongly amplified (Ct value < 30) the relevant target genes for all 12 control panel isolates listed in Table 1, allowing each isolate to be classified into the appropriate clade (Table 4). There was no weak positive (Ct values > 30) or off-target amplification of the target genes was observed for the Regional and Clade real-time PCR assays (Table 4). Using a dilution series, the limit of detection was determined as the lowest DNA concentration resulting in a true positive (Ct < 30). The limit of detection for these assays was determined to be 0.1 μM (Table 6).

**Table 4.** Average cycle threshold (Ct) values from the clade-typing real-time PCR assays performed using the control panel isolates. Ct values under 30 indicate a positive result and Ct values over 30 indicate a negative result (shown as “-“)

| Real-time PCR Ct values for target genes |  |  |  |  |  |  |
| --- | --- | --- | --- | --- | --- | --- |
| Isolate name | Expected Clade* | *BTN76_08545* | *SEN1975* | *pemI* | *SEN1943* | Real-time PCR clade result |
| P125109 | Global Epidemic | - | 19.31 | - | 17.72 | Global Epidemic |
| A1636 | Global Epidemic | - | 19.26 | - | 18.12 | Global Epidemic |
| 1320 | Global Epidemic | - | 18.54 | - | 16.67 | Global Epidemic |
| 791 | Global Outlier | - | 19.9 | - | - | Global Outlier |
| 672246 | Global Outlier | - | 17.67 | - | - | Global Outlier |
| 672632 | Global Outlier | - | 18.23 | - | - | Global Outlier |
| D7795 | East African | 17.66 | - | 16.92 | - | East African |
| CP225 | East African | 18.12 | - | 16.81 | - | East African |
| 6396 | East African | 17.05 | - | 16.6 | - | East African |
| 10136/01 | West African | 18.59 | - | - | - | West African |
| 0527/01 | West African | 18.74 | - | - | - | West African |
| 8078/01 | West African | 20.26 | - | - | - | West African |

\*Derived from Feasey *et al* 2016 clade-typing using whole genome sequences

**Table 5.** Efficiency of multiplex assays based on the average Ct values and performance analysis of assays for each primer and probe set performed with three technical replicates

| DNA concentration<br>(ng/ μL) | African Cluster<br>target* | Global Cluster<br>target* | East African Clade<br>target* | Global Clade<br>target* |
| --- | --- | --- | --- | --- |
| 10 | 23.33 ± 0.34 | 21.95 ± 0.49 | 22.73 ± 0.48 | 21.36 ± 0.26 |
| 1 | 27.34 ± 0.08 | 25.55 ± 0.81 | 26.08 ± 0.28 | 23.99 ± 0.54 |
| 0.1 | 28.89 ± 0.59 | 28.96 ± 0.39 | 27.97 ± 0.42 | 27.15 ± 0.49 |
| 0.01 | 34.00 ± 0.54 | 32.12 ± 0.46 | 32.62 ± 0.12 | 30.05 ± 0.48 |
| 0.001 | 35.93 ± 0.65 | 35.15 ± 0.90 | 35.30 ± 0.09 | 34.12 ± 0.49 |
| **Determination coefficient ($R^2$) | 0.98 (0.95 - 1.00) | 0.99 (0.98 - 1.01) | 0.99 (0.98 - 1.00) | 0.99 (0.98 - 0.99) |
| Slope | 0.30 | 0.30 | 0.31 | 0.31 |
| Slope derived efficiency | 1.00 | 1.00 | 1.04 | 1.04 |

\*Average Ct values ± standard deviation (n = 3) for 5 target DNA concentrations
\*\* Determination coefficient (numbers in brackets indicate 95% Confidence Interval)

**Table 6.**
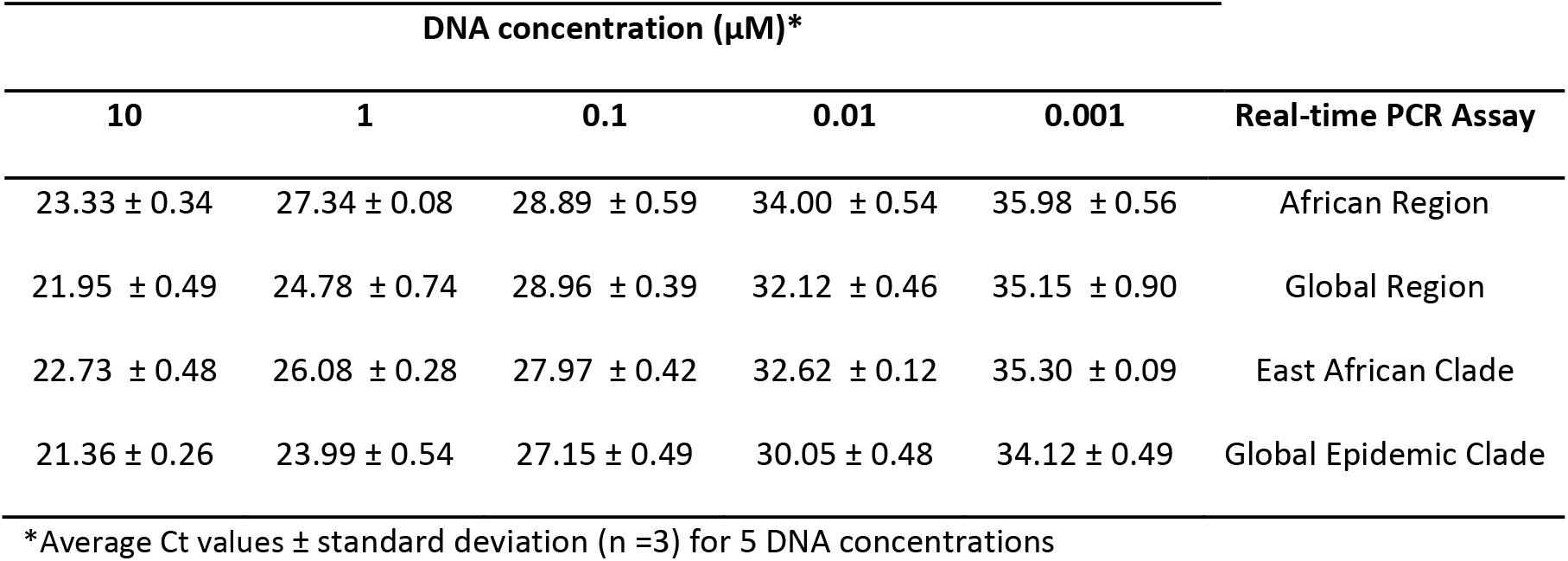
Cycle threshold (Ct) values from clade-typing real-time PCR assays performed with a DNA dilution series

| DNA concentration (μM)* | | | | | |
| --- | --- | --- | --- | --- | --- |
| 10 | 1 | 0.1 | 0.01 | 0.001 | Real-time PCR Assay |
| 23.33 ± 0.34 | 27.34 ± 0.08 | 28.89 ± 0.59 | 34.00 ± 0.54 | 35.98 ± 0.56 | African Region |
| 21.95 ± 0.49 | 24.78 ± 0.74 | 28.96 ± 0.39 | 32.12 ± 0.46 | 35.15 ± 0.90 | Global Region |
| 22.73 ± 0.48 | 26.08 ± 0.28 | 27.97 ± 0.42 | 32.62 ± 0.12 | 35.30 ± 0.09 | East African Clade |
| 21.36 ± 0.26 | 23.99 ± 0.54 | 27.15 ± 0.49 | 30.05 ± 0.48 | 34.12 ± 0.49 | Global Epidemic Clade |

\*Average Ct values ± standard deviation (n =3) for 5 DNA concentrations

### Performance analysis of the real-time PCR assays

To determine assay efficiency, the Region and Clade real-time PCR assays were performed using serial dilutions (10-fold) of the genomic DNA extracted from two control isolates (D7795 and A1636), and calibration curves were plotted to assess the linear range (assessment of how well the assay amplifies the target gene at various DNA concentrations (R^2^)) and the amplification efficiency (how well the assay amplifies the target gene region).

The Region real-time PCR assay that contained African and Global Region primer and probe sets had linear ranges of 0.98 and 1.00 respectively (Table 5). The Clade real-time PCR assay that contained East African clade and Global Epidemic clade primer and probe sets both had linear ranges of 0.99 (Table 5). Thus, the linear range for the clade-typing assay complied with the required R^2^ ≥ 0.98 (25), meaning that the primer and probes for the Region and Clade real-time PCR assays efficiently amplified the target genes. The amplification efficiencies were calculated based on the slope of calibration curves. The theoretical maximum amplification efficiency is 1.00, which indicates that the amount of product doubles with each cycle (26). The Region and Clade assays performed at an average efficiency of 1.00 and 1.04 respectively (Table 5).

## 7. Conclusion

Here, we have described the development of a high-quality molecular classification assay for the clade-typing of *S*. Enteritidis that is ideal for use in public health laboratories, especially where WGS is not readily available. All primer and probe sets for the Region and Clade assays ran at optimal efficiency within the multiplex assays. This novel multiplex PCR assay could be used to investigate whether certain clades of *S*. Enteritidis because human disease of differing severity.

## 8. Author statements

### 8.1 Authors and contributors

Sarah Gallichan: Investigation; Experimental Design; Validation of Methodology

Nicholas Feasey: Supervision; Review & editing

Blanca Perez-Sepulveda: Resources; Review & editing; Validation of Methodology

Jay Hinton: Resources; Review & editing

Juno Thomas: Supervision

Anthony Marius Smith: Supervision; Review & editing

### 8.4 Ethical approval

Ethical clearance for all laboratory-based surveillance and research (approved 12 November 2018) was obtained from the University of Witwaterstrand, Johannesburg, South Africa (Wits protocol no. M140159) by the Centre for Enteric Diseases, National Institute for Communicable Diseases.

